# Detection of native and mirror protein structures based on Ramachandran plot analysis by interpretable machine learning models

**DOI:** 10.1101/2020.09.03.280701

**Authors:** Julia Abel, Marika Kaden, Katrin Sophie Bohnsack, Mirko Weber, Christoph Leberecht, Thomas Villmann

**Author notes:** These authors contributed equally to this work (Julia Abel, Marika Kaden, Thomas Villmann). These authors also contributed equally to this work (Katrin Sophie Bohnsack, Mirko Weber, Christoph Leberecht).

## Abstract

In this contribution the discrimination between native and mirror models of proteins according to their chirality is tackled based on the structural protein information. This information is contained in the Ramachandran plots of the protein models. We provide an approach to classify those plots by means of an interpretable machine learning classifier - the Generalized Matrix Learning Vector Quantizer. Applying this tool, we are able to distinguish with high accuracy between mirror and native structures just evaluating the Ramachandran plots. The classifier model provides additional information regarding the importance of regions, e.g. *α*-helices and *β*-strands, to discriminate the structures precisely. This importance weighting differs for several considered protein classes.

## Introduction

Proteins can have two conformations: the L-enantiomeric conformation and the D-enantiomeric conformation. The first represents the natural form (further known as native) of a protein, whereas the latter represents an exact mirror-image of it [21, 52, 60, 63]. The chemical general term for two enantiomeric conformations is chirality. By definition these enantiomers are mirror-images of each other and not superimposable [62].

Although the two enantiomeric forms of one protein exhibit more or less the same behavior concerning physical and chemical properties as well as substrate specificity [39, 52, 63], mirror proteins can show a more stable behavior towards natural enzyme degradation and reduced immunogenicity [21, 41, 60].

The differentiation of native and mirror proteins is crucial for further analysis and research in the fields of drug discovery/development and synthetic biology. [59, 61]. Especially for the latter holds that mirror proteins give the opportunity to create a fully working molecular system, which would not be dependent on natural compounds [60] and, therefore, orthogonal systems would be possible [30]. Furthermore, mirror proteins hold the possibility of creating mirror life [30]. The presence of native and mirror models is of particular importance in the structure reconstruction of proteins from amino acid sequences. Contact maps can be used as a set of constraints for *ab initio* structure prediction. They show predicted contacts based on co-evolving residues which imply their spatial proximity and are derived from a multiple sequence alignment [6]. Contact map based approaches may lead to either native or mirror models since a contact map contains no information regarding chirality and both, native and mirror proteins exhibit the same inter-atomic distances [27]. In order to screen out such mirror models, a variety of approaches were published, which either intervene in the process of model generation [16, 32] or filter mirror models subsequently [11, 55].

Kurczynska and Kotulska tried in their work [28] to distinguish between native and mirror models in an automated way (Automated Mirror-Native-Discrimination – AMND). For this purpose, they used the energy terms obtained by PyRosetta, a Python-based interface to Rosetta for molecular modeling [9]. The discrimination results of 69% in average are respectable but far away to be satisfying. Further, it was not possible for them to find one uniform rule based on that energy terms for all the proteins classes **A**-**G** of SCOPe [18, 31], which were studied in this investigation. Albeit in [2] it becomes apparent that the calculation of the energy terms is rather complex. Alternatively, they also took into account to consider the differences within Ramachandran plots (R-plots), and thus the dihedral angles distribution of a protein’s backbone by depicting them in a toroidal plot, because these plots are known to provide structural information [43]. However, they explained that these plots are not feasible for automated mirror/native discrimination.

Nonetheless, there are three things a R-plot definitely can give information about: a protein’s secondary structure elements, the favored/allowed regions of those and also the handedness of helices (right-handed and left-handed) [3, 48]. Following the International Union of Pure and Applied Chemistry’s (IUPAC) definition of handedness being the same as chirality [37], the distinctive feature for differentiating between native and mirror models would be the handedness of helices.

Since this handedness can also be found as a characteristic of R-plots, they represent a simple but important tool for finding structural differences in proteins [3]. Assuming that native and mirror proteins display different structures the resulting differences in the R-plots should be observable.

To tackle the AMND problem, appropriate mathematical and statistical tools are required, which reflect the mathematical structure of the R-plots and can evaluate them in an automatic way with sufficient certainty. Therefore machine learning tools like (deep) artificial neural networks come into play [13], which are powerful tools to analyze complex data like R-plots. However, these networks and their inferences are frequently not explainable [45]. An alternative to deep networks provide interpretable machine learning approaches like prototype based models [5, 56], which rely on the nearest prototype principle for data representation [38]. Frequently, these networks make use of specific data properties [58] and, hence, their results are better to interpret and may provide additional knowledge drawn from the training data.

In this context, a key observation regarding R-plots is that we can take them as approximated density plots or histograms of dihedral angle pairs (Φ, Ψ) in the two-dimensional plane. Thus, the data analysis and statistical evaluation have to deal with the comparison of densities and their discrete representations. Thus, we apply an adaptive vector quantization based classifier model – the so-called Generalized Matrix Learning Vector Quantizer (GMLVQ) [19, 50]. After training this classifier provides structural knowledge regarding the data, which supports the classification decision. In particular, a classification correlation matrix is delivered, which describes correlations between data [5, 56]. GMLVQ is known to be robust and easy to interpret [46, 58]. In biomedical context it was successfully applied to analyse flow cytometry data and to detect early folding residues during protein folding [4, 7].

The structure of this paper is as follows: First the data set in use is described more detailed as well as the corresponding data preprocessing. Afterwards we give a brief introduction of R-plots as well as the machine learning data analysis tool GMLVQ, which is an interpretable artificial neural network. Subsequently, we state the general workflow for the AMND to distinguish between native and mirror samples based on R-plot analysis by means of GMLVQ.

Finally we present the numerical results for classification performance and the extracted knowledge provided by the interpretable model together with its direct biological explanation.

## Materials and methods

### Ramachandran Plots

*Ramachandran plots* (R-plots) display the dihedral angles Φ and Ψ of a protein’s backbone to visualize their distribution [43]. R-plots provide an easily inspectable tool to detect underlying properties of the secondary structure in that protein [3]. However it has to be kept in mind, that the measured angles describe the current state at the time of measurement of the respective dynamic atoms in the backbone [33]. Mathematically, R-plots can be taken as approximated probability distributions for pairs (Φ, Ψ) of dihedral angles. Accordingly, if we partition the plot by an *N* × *N* cell array as depicted in S1 Fig, we can describe each plot as a vector **x** = (*x*_1_, …, *x*_*n*_)^*T*^ ∈ ℝ^*n*^ with *n* = *N* ^2^, where *x*_*k*_ ≥ 0 is the (relative) number of dihedral pairs in the cell with the coordinates (*j, l*) where *k* = (*j* − 1) *· N* +*l*. Thus, **x** is a relative histogram vector also denoted as probability vector, i.e. 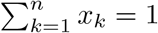 S1 Fig shows a generic R-plot of pairs (Φ, Ψ), for a backbone. The horizontal axis is related to Φ, whereas the vertical axis is related to Ψ. The allowed/favored areas of secondary structures are shown. Particularly, we indicate the regions of *β*-sheets (*β*-strands), as well as right- and left-handed *α*-helices. Additionally we have added a 6 × 6 cell grid to the R-plot, so that we can specifically address cells of the R-plot by their coordinates.

For example cells (1,4) and (2,4) as well as cells (4,1) and (4,2) characterize the favored regions for right-handed *α*-helices (*α*) and left-handed *α*-helices (**L**_*α*_), respectively. The cells (1,2) and (2,1) constitute the favored regions for *β*-strands (*β*).

### The Generalized Learning Vector Quantizer – An Adaptive, Interpretable Classifier in Machine Learning

*The Generalized Learning Vector Quantizer* (GLVQ, [47]) is a prototype-based machine learning method for classification derived from a heuristic approach proposed by T. Kohonen [22]. The mathematically well justified model optimizes the classification performance by means of stochastic gradient descent learning (SGDL, [14, 44]) and is known to be robust and interpretable. The underlying cost function to be minimized during training approximates the overall classification error for a given training data set *T* = *{*(**x**_*j*_, *c* (**x**_*j*_)) ∈ *X* × 𝓁, *j* = 1 …*N }*, where *X* ⊆ ℝ^*n*^ is the set of training vectors and *c* (**x**_*j*_) ∈ 𝓁 = {1, …, *C*} is a training data class label out of a set of available class labels [19]. For this purpose, the GLVQ model assumes a set of prototypes *W* = *{***w**_*k*_ ∈ ℝ^*n*^, *k* = 1 …*M }* with class labels *c* (**w**_*k*_) ∈ 𝓁 such that at least one prototype is assigned to each class. Thus, a partition of the prototype set 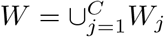 with *W*_*j*_ = *{***w**_*k*_ ∈ *W* |*c* (**w**_*k*_) = *j}* is obtained, i.e. *W*_*j*_ ≠ ∅ for all *j* = 1 …*C* and *W*_*i*_ ∩ *W*_*j*_ for *i* ≠ *j* are valid.

Further, a dissimilarity measure *d* (**x, w**) is supposed to evaluate the similarity between data and prototypes. This measure is required to be differentiable at least with respect to the second argument to ensure SGDL.

For a given GLVQ-configuration, i.e. a fixed prototype set *W*, a new data point **x** is classified by the assignment

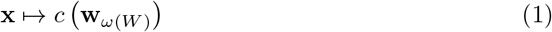

with

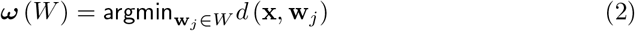

known as winner-takes-all in nearest prototype classification [38]. The prototype **w**_*ω*_ is denoted as winner of the competition.

During the learning, GLVQ adapts the prototypes minimizing the overall cost function

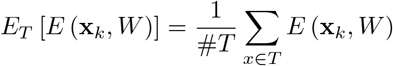

by SGDL with respect to the prototypes realizing a competitive learning scheme [1], where

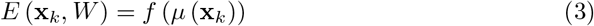

is the local classification error and #*T* denotes the cardinality of the training data set. The local error *E* (**x**_*k*_, *W*) depends on the choice of the monotonically increasing squashing function *f* and the classifier function

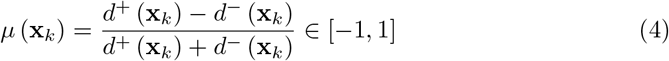

where *d*^*±*^ (**x**_*k*_) = *d*^*±*^ (**x**_*k*_, **w**^*±*^) and 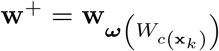 is the so-called best matching correct prototype and 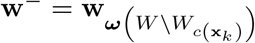 is the corresponding best matching incorrect prototype. Frequently, the squashing function *f* is chosen as a parametrized sigmoid: 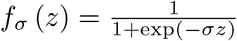 with the slope parameter *σ* [57].

The classifier function *µ* (**x**_*k*_) becomes negative for correctly classified training data. In fact, the cost function *E*_*T*_ [*E* (**x**_*k*_, *W*)] is an approximation of the overall classification error. The approximation precision depends on the specific choice of the squashing function and their parameters. However, in case of the mentioned parametrized sigmoid function, the GLVQ is robust regarding moderate variations of the steepness parameter. For a detailed consideration we refer to [20].

In standard GLVQ, the squared Euclidean distance *d*_*E*_ (**x, w**) = (**x** − **w**)^2^ is used. Another popular choice is the squared *Euclidean mapping distance*

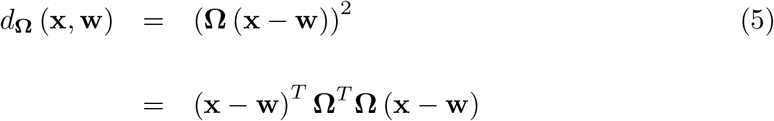

proposed in [50] depending on the mapping matrix **Ω** ∈ ℝ^*m*×*n*^. Here, *m* is the mapping dimension usually chosen as *m* ≤ *n* [8]. Thus, the data are first mapped linearly by **Ω** and afterwards the Euclidean distance is calculated in the mapping space ℝ^*m*^.

The SGDL for training performs an attraction-repelling scheme in the data space as the common feature for all LVQ-schemes [23]: For a given training data **x**_*j*_ with known class label *c* (**x**_*j*_), the best matching correct prototype **w**^+^ is moved a trifle towards **x**_*j*_ whereas the best matching incorrect prototype **w**^−^ is slightly repelled. Iterative application of that scheme with random training data selection realizes SGDL for the cost function *E*_*T*_ [*E* (**x**_*k*_, *W*)].

Interestingly, the mapping matrix **Ω** can also be optimized by SGDL to achieve a better separation of the classes in the mapping space [51]. Note that SGDL for **Ω**-optimization usually requires a careful regularization technique [49]. The respective algorithm is known as *Generalized Matrix LVQ* (GMLVQ) [50].

After training, the adjusted mapping matrix **Ω** provides additional information about the data features with respect to the classification task. More precisely, the resulting matrix **Λ** = **Ω**^*T*^ **Ω** ∈ ℝ^*n*×*n*^ allows an interpretation as *classification correlation matrix*, i.e. the matrix entries Λ_*ij*_ reflect the strength of the correlations between the data features *i* and *j* contributing to the class discrimination [5, 56]. The diagonal entries Λ_*ii*_, which are all non-negative, are the relevances *r*_*i*_ = Λ_*ii*_ delivering the importance of the *i*th data feature for class separation: The higher the relevance value *r*_*i*_ the more important is the corresponding feature. Relevance values *r*_*i*_ ≊ 0 indicate that the respective features are not important for class separation or that the information is already provided by other features [15]. All relevance values *r*_*i*_ are collected in the relevance profile vector **r** = (*v*_1_, …, *v*_*n*_)^*T*^. The process of relevance vector adaptation is also denoted as *relevance learning*.

Further, this approach allows to detect outliers due to the bounded minimum distance principle in contrast to class assignments via decision hyperplanes [12]. Accordingly, outliers are rejected because the model validity cannot guaranteed for these data. (*outlier-reject*). Further, data **x** with uncertain classification decision can be rejected evaluating the difference between the distance *d* (**x, w**_*ω*(*W*)_) to the overall winner **w**_*ω*(*W*)._ and the distance *d* (**x**, *ω* (*W*^*^)) with 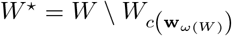. In case of a significant deviation, which corresponds to an uncertain decision, the query regarding **x** is rejected (*classification-reject*).

### The Data Set

We used exactly the same data set as given in [28], which was generated and analyzed there to detect native and mirror structures. Therefore, we give only a short summary of the data generation and preprocessing, for a more detailed description it is referred to [28]:

In order to emulate the necessary procedures in structure modeling from contact maps, the authors derived these maps [17] from a set of 1, 305 representative domains of SCOPe superfamilies. Those maps were in turn the base for reconstructing full-atom proteins of different configurations [24, 25, 29, 54], approximately 50 native and 50 mirror models for each domain. Consequently the whole data set comprises 130, 500 models, from which the dihedral angles were calculated [10].

Further all of the models in the data set can be grouped into one of the classes **A**-**G** of SCOPe, representing seven distinctive data sets. Class **A** (*all α*) describes proteins that are predominantly made of *α*-helices, whereas class **B** (*all β*) mainly consists of *β*-strands. Proteins with alternations of *α*-helices and *β*-strands are put into class **C**. In contrast, proteins in class **D** show segregated sections of *α*-helices and *β*-strands. So this four classes are categorized due to secondary structure features of the proteins, whereas the following three are not. The so-called multi-domain proteins are grouped into class **E**. Proteins and peptides located on surfaces or in membranes are grouped into class **F**. Small proteins, that have little or no secondary structure, are grouped into class **G** [18, 28, 31, 48]. In Tab. 1 the sizes of these groups are depicted. All of these models were condensed into the class **ALL**.

**Table 1.** Numbers of samples for each of the described protein classes (domains) **A − G**

| protein class | protein models |
| --- | --- |
| **A** | 343 |
| **B** | 233 |
| **C** | 149 |
| **D** | 368 |
| **E** | 21 |
| **F** | 78 |
| **G** | 113 |

### Data Preprocessing and Machine Learning Settings

Each of the aforementioned data sets determines a specific learning task to distinguish mirror and native samples. To do so a relative R-plot histogram vector **x** ∈ ℝ^*n*^ with *n* = *N* × *N* with *N* = 6 as grid resolution was extracted for each sample. These data vectors served as training data for GMLVQ according to the considered tasks.

For each learning task **A**-**G**, and **ALL** we trained a separate GMLVQ model with three prototypes per class (mirror/native). Additionally to the prototypes we also adapted the mapping matrix **Ω** with the mapping dimension *m* = 36. The reported classification results are achieved as the averaged test performances obtained by 50 independent runs. Each run was done as a five-fold cross-validation procedure.

Further, for visual inspection and evaluation, we generated a *summarized R-plot*, which is collecting all pairs (Φ, Ψ) of dihedral angles for all samples of the considered task in a single R-plot but separately for native and mirror samples. Thus, these R-plots can be seen as estimated dihedral angles densities in the (Φ, Ψ)-plane for mirror and native samples, allowing a visual inspection of those (Φ, Ψ)-distributions, see S2 Fig–S4 Fig.

## Results and Discussion

By taking advantage of the inherent interpretable nature of GMLVQ, particularly by considering the relevances of the data features, we can extract insightful knowledge regarding the classification decision. Especially the connection of the relevant features with the favored and allowed regions in the R-plots for each protein class offers a biological interpretation. Tab. 2 summarizes the areas of the R-Plot of each protein class which are considered most relevant by the model.

**Table 2.**
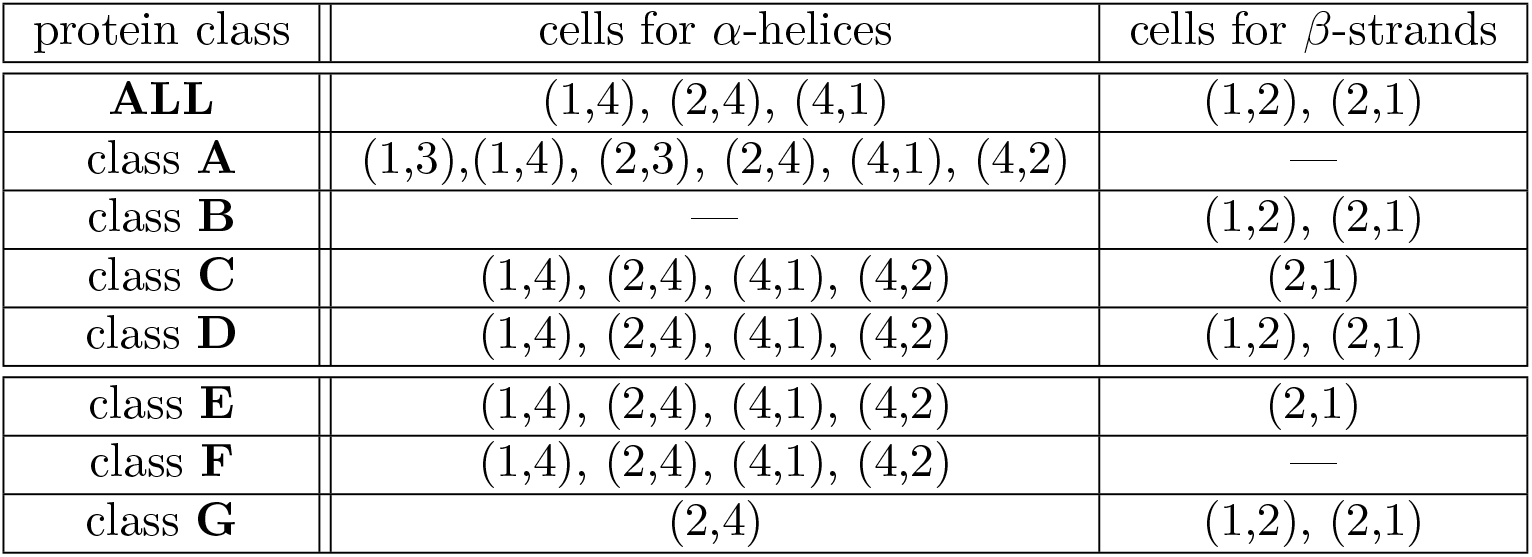
Most relevant cells for differentiation of mirror and native models based on the corresponding relevance profiles.

As previous publications suggest [28, 55], native and mirror conformations of proteins rich in helices (belonging to class **A**), can be distinguished based on chirality derived information, or in more detail: right-handed *α*-helices are favoured in native conformations [40]. We can confirm this finding, since the features corresponding to the left- and right-handed *α*-helices predominantly contribute to the class discrimination of mirror and native (see S3 Fig and Tab. 2). The accuracy for this class is 86.57% (see Tab. 3 for all accuracies). Interestingly, our model achieves an accuracy of 92.56% for class **B**, which obviously collides with the statement of those native and mirror models being indistinguishable [28, 55] and, furthermore, even exceeds the accuracy of class **A**. The relevant features for class discrimination are those corresponding to *β*-strands in the R-plot (see S3 Fig). In detail, the important underlying secondary structures in this case might be the right-handed triple helices (collagen) and parallel *β*-strands [26]. However, the confirmation of the actual underlying secondary structure as well as its relation to chirality are still pending.

**Table 3.**
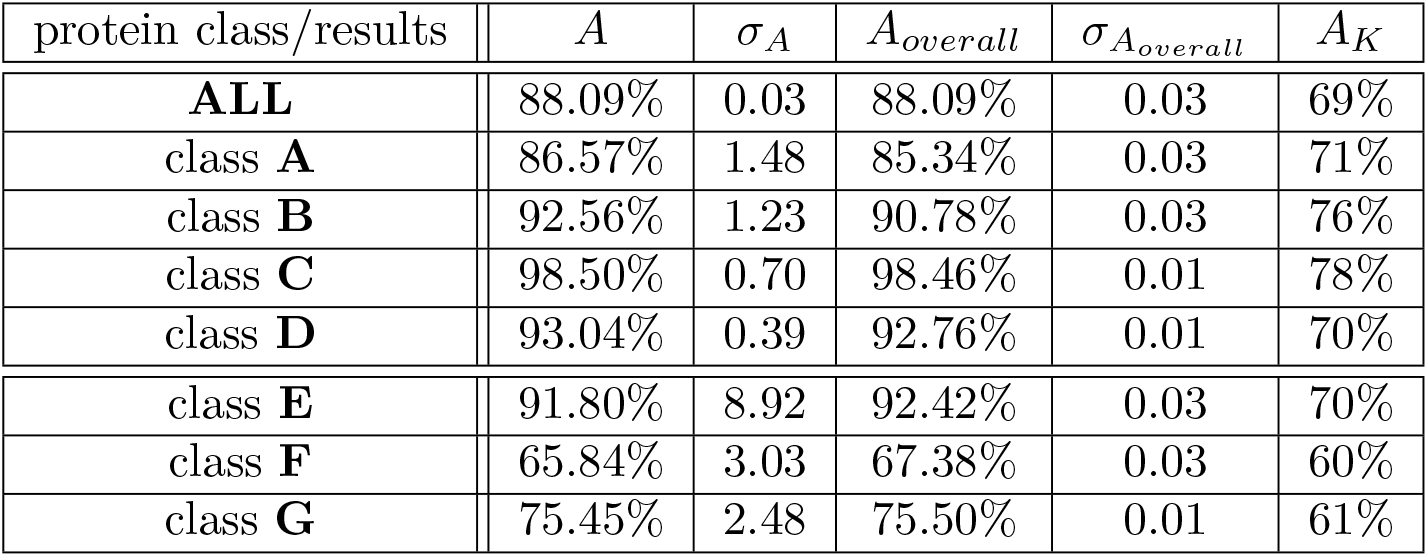
Obtained averaged (test) accuracies (A), obtained accuracies using the general model for specific classes together with their respective standard deviations for the protein classes (results refer to 50 runs of cross-validated GMLVQ). For comparison, additionally the best classification results *A*_*K*_ from [28] are given (for all classes as average of the others)

As protein classes **C** and **D** structurally show a combination of the aforementioned two classes *all-alpha* and *all-beta*, the relevant features for class discrimination also do. Class **C** shows the best of all investigated accuracies. This result concurs with the findings in [28]. Among the protein classes which were not categorized due to their secondary structure, the multi-domain class **E** by far shows the best accuracy with 91.8%, whereas classes **F** and **G** do not exceed 80%. Even though class **E** has got such a high accuracy it has to be treated with care, since there has not been enough data for this protein class. The relatively poor accuracy for class **F** is most likely due to the fact that membrane proteins propose some difficulties in structure elucidation [34, 42, 64] and this results in low resolutions [36]. A poor resolution in turn may lead to inaccurate atom coordinates. That means, the calculations of the dihedral angles cannot be correct either and, therefore, complicate the classification. As for class **G** the obtained low accuracy is probably due to the fact that small proteins do not have that many amino acids, less than 100 [35, 53], and therefore show less *α*-helices or *β*-strands than other proteins.

The R-plots for classes **E**–**G** together with the corresponding relevance profiles are depicted in S3 Fig.

In order to assess the models suitability for a more general problem we considered all protein classes for training as well as for testing and achieved an overall accuracy of 88.09%. Pursuing this approach, which is more general and more considerable for application, we investigated the behavior of our model by training with all protein classes but testing with only one protein class at a time. The achieved accuracies are in good agreement with those of the single classes as Tab. 3 shows.

## Conclusion

In the present contribution we offer a valid approach for distinguishing mirror and native conformations of proteins based on structure information. The approach is based on the evaluation of the respective R-plots by means of an interpretable machine learning model. This model, the Generalized Learning Matrix Vector Quantizer, is known to be robust and highly interpretable according to the underlying reference principle. Moreover, according to the integrated relevance learning metric adaptation, the approach provides beside the classification ability additional knowledge regarding the classification decision. In the context of R-plot analysis this information consists in a weighting of importance of R-plot regions regarding best mirror-native-separability.

Kurczynska and Kotulska state that structural features are no discriminatory property of native and mirror models. Although information regarding chirality can be used to differentiate models rich in *α*-helices, according to Vendruscolo et al. this does not hold for all-*β* structures [55]. However, we were able to show that a discrimination of native and mirror models using structural features is indeed possible.

The GMLVQ classifier achieves high separation accuracies for all protein classes except class **F** and **G**. At least for the latter one, acceptable results are obtained. In fact, the resulted accuracies for protein classes **F** – **G** show that a distinction of mirror and native structures by means of R-plots is possible with high precision and sensitivity. The interpretable model offers additional insights: In particular, the relevance profiles, weighting the regions like *α*-helices and *β*-strands of R-plots for mirror-native-discrimination, differ for the considered protein classes. The obtained relevance profiles are in good agreement with respective biological knowledge about protein structure chirality. at least for the considered data set.

Thus, the presented approach offers a successful alternative to the statistical approach based on energy levels as proposed in [28] and emphasize the importance of R-plots for structural analysis of proteins as already mentioned in [3]. Along this line, also data processing is easier in the present approach compared to the complex calculations of the energy levels [2].

Further investigation should include to improve the classification performance by finer cell resolutions of R-plots. However, this requires more training data for sufficient learning stability. Further, reject option strategies should be included to detect outliers.

## Acknowledgments

The authors greatly acknowledge helpful discussions within the CI-research group of the SICIM at the UAS Mittweida.

## Supporting Information

J.A., K.S.B., and M.K. were supported by a grant for a so-called *Young Researcher Group* provided by the European Social Fund (ESF).

**S1 Fig.**
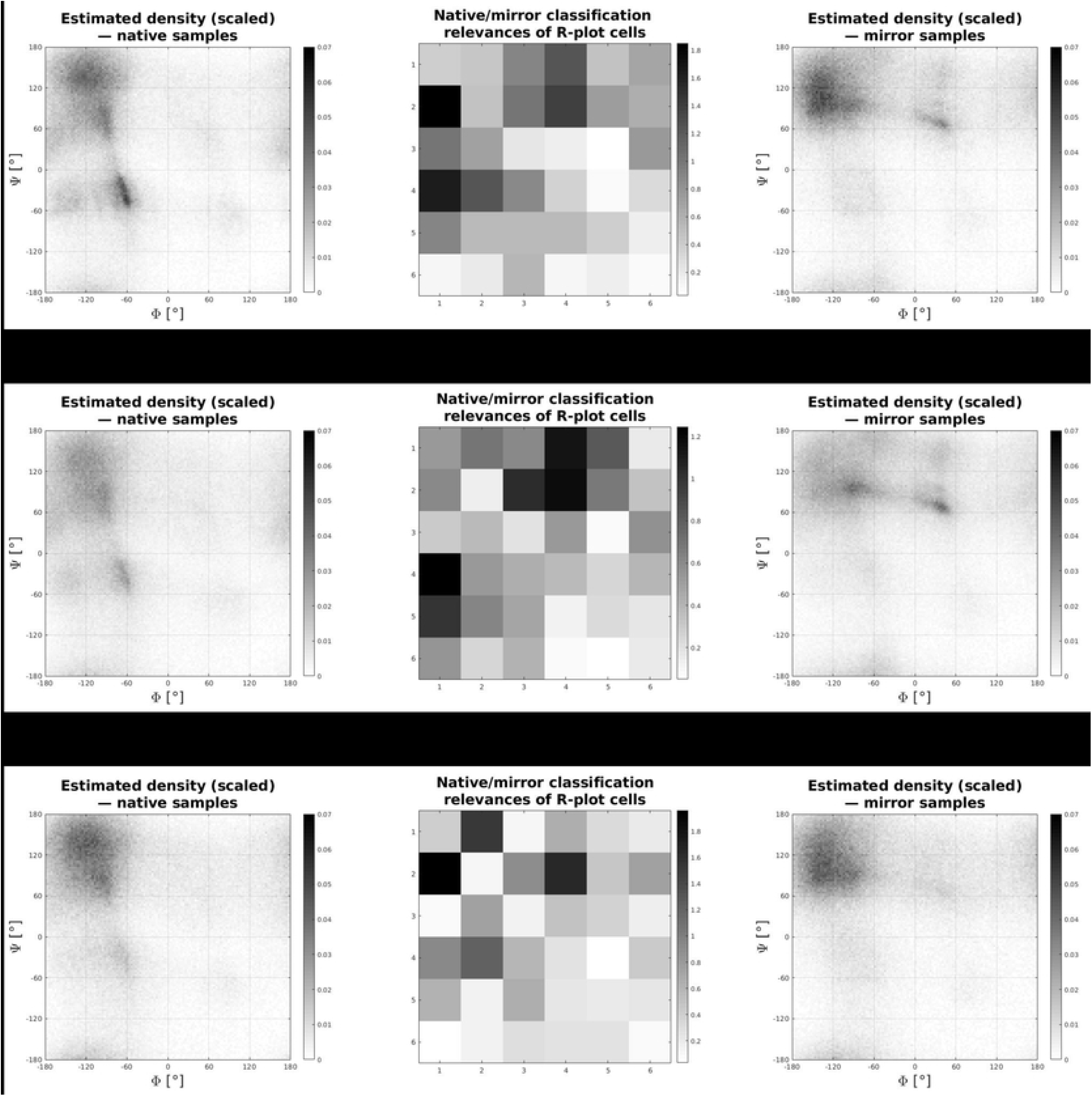
Generic Ramachandran plot (R-plot) for dihedral angles. The allowed/favored regions of secondary structures are shown. Particularly, we indicate the areas of *β*-sheets/-strands (*β*), as well as right-handed *α*-helices (*α*) and left-handed *α*-helices (L_*α*_). Additionally we have added a 6 × 6 cell array to the R-plot, so that we can address regions (cells) of the R-plot accordingly.

**S2 Fig.**
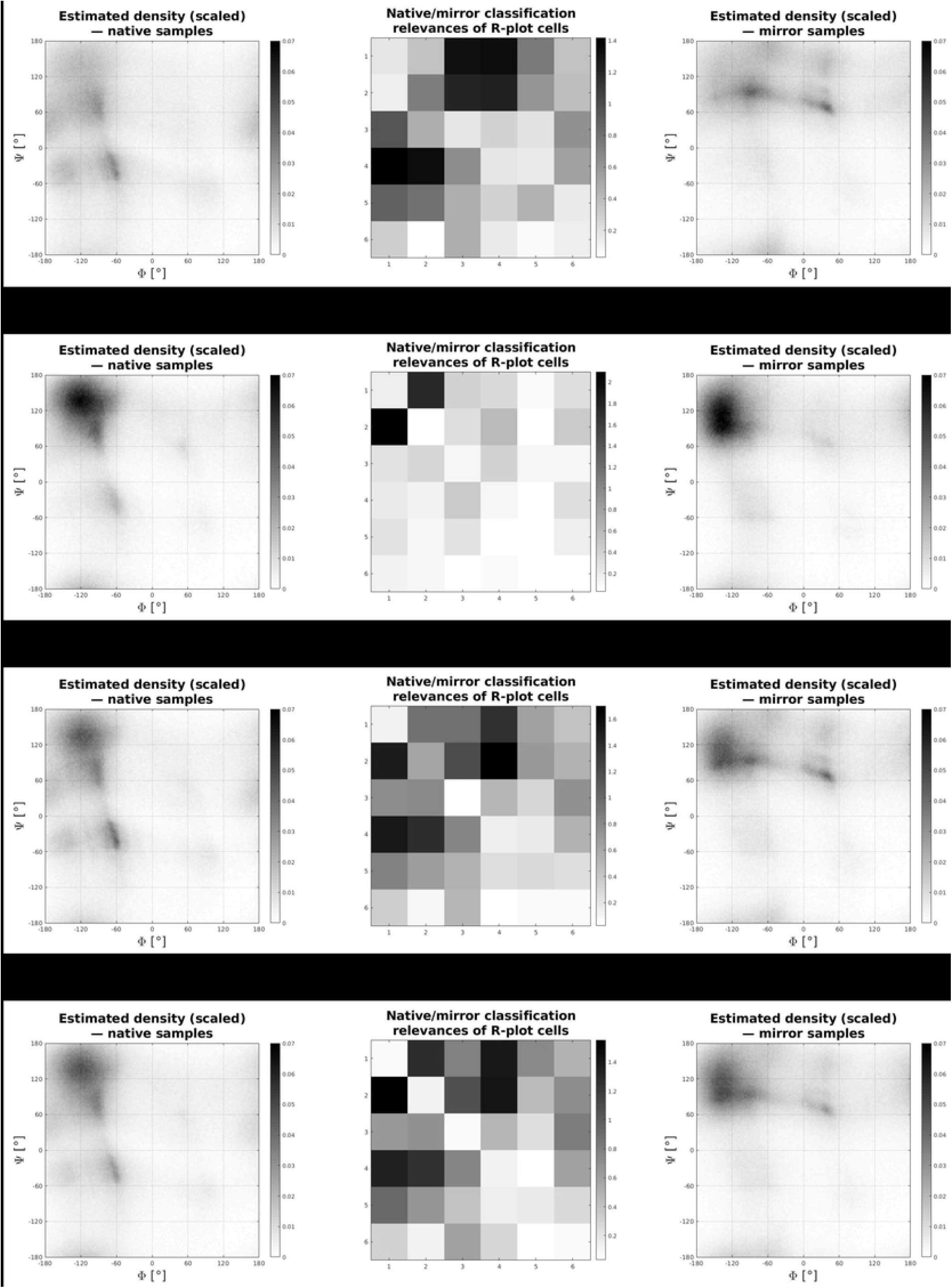
Summarized R-plots and respective cell relevance for mirror/native models for learning of all data (ALL). The summarized R-plots for native (left) and mirror (right) samples. The plots are estimators for the dihedral angles densities in the (Φ, Ψ)-plane. The relevance profile vector **r** = (*r*_1_, …, *r*_36_)^*T*^ is depicted in the middle, arranged accordingly to the cells. The higher (darker) the *r*_*i*_-value the more important is the respective cell for AMND. The achieved GMLVQ classification accuracy for this data set is 88.09%.

**S3 Fig.**
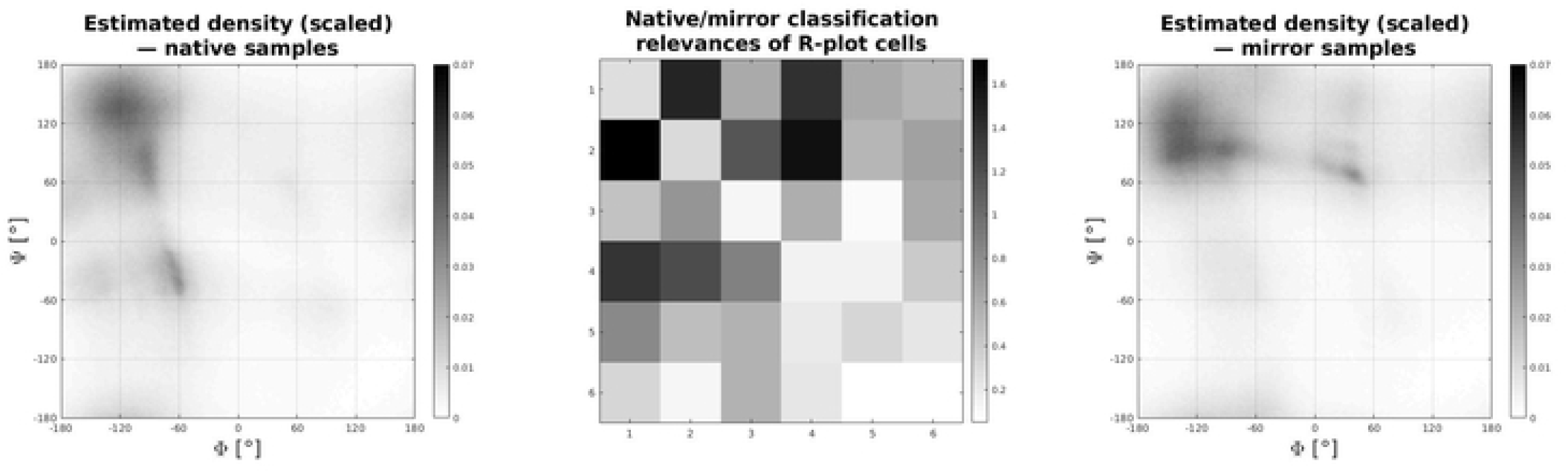
Summarized R-plots and respective cell relevance for mirror/native models for learning of the classes A–D. The summarized R-plots for native (left) and mirror (right) samples. The plots are estimators for the dihedral angles densities in the (Φ, Ψ)-plane for samples. The obtained relevance profile vectors **r** = (*r*_1_, …, *r*_36_)^*T*^ from relevance learning in GMLVQ are depicted in the middle, arranged accordingly to the cells of the R-plot. The higher (darker) the *r*_*i*_-value the more important is the respective cell for AMND. The achieved GMLVQ classification accuracies for these data sets are given in Tab. 3.

**S4 Fig.**
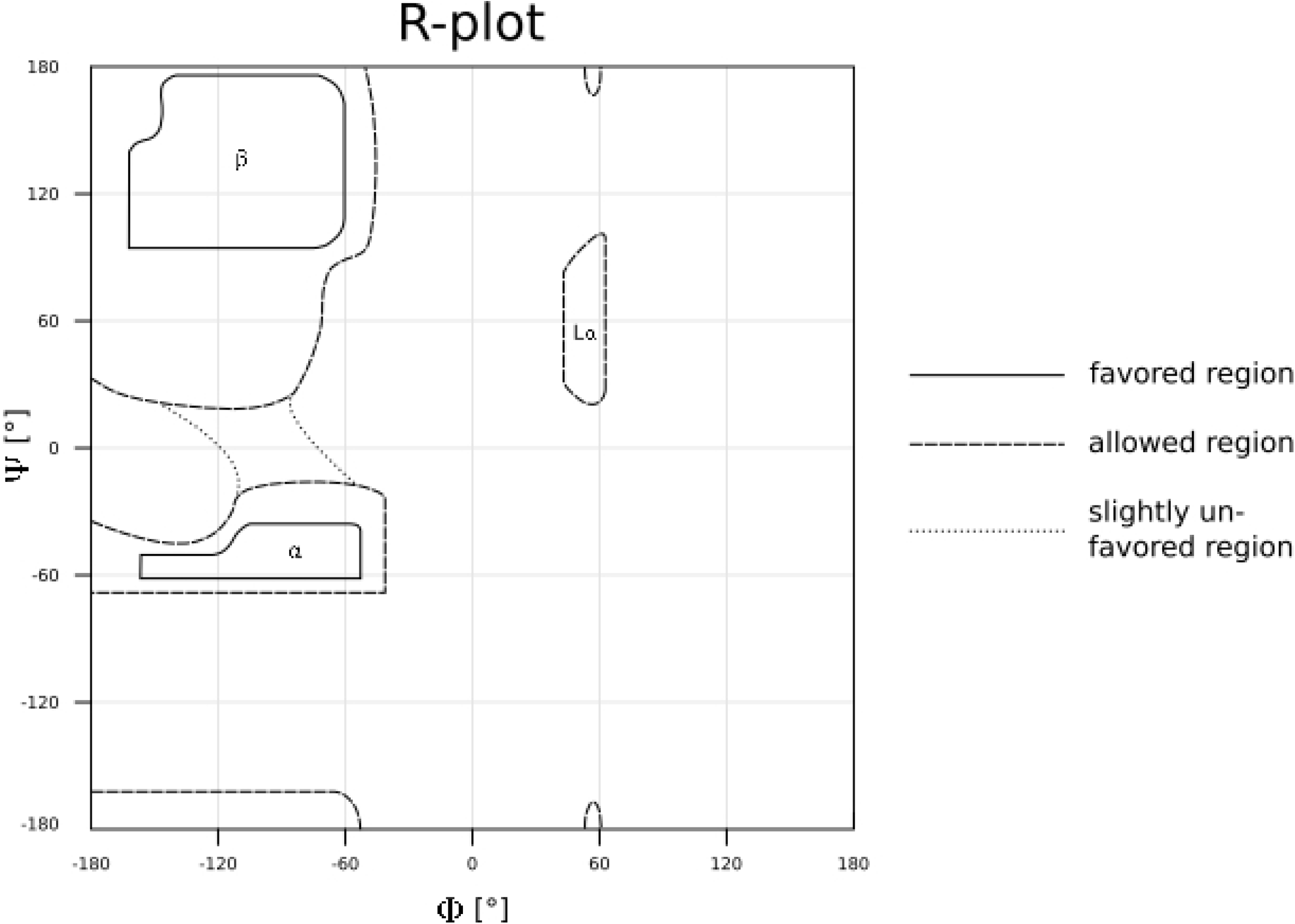
Summarized R-plots and respective cell relevance for mirror/native models for learning of the class E–F. The summarized R-plots for native (left) and mirror (right) samples. The plots are estimators for the dihedral angles densities in the (Φ, Ψ)-plane for samples. The obtained relevance profile vectors **r** = (*r*_1_, …, *r*_36_)^*T*^ from relevance learning in GMLVQ are depicted in the middle, arranged accordingly to the cells of the R-plot. The higher (darker) the *r*_*i*_-value the more important is the respective cell for AMND. The achieved GMLVQ classification accuracies for these data sets are given in Tab. 3.

**S5 Tab**. As an add-on we provide in Tab. 4 the sensitivities as well as the specifitities corresponding to the reported accuracies from Tab. 3.

**Table 4.** Overall model (test) sensitivities (*Se*) and specificities (*Sp*) together with their respective standard deviations for the protein classes (results refer to 50 runs of cross-validated GMLVQ)

| protein class / results | $Se$ | $\sigma_{Se}$ | $Sp$ | $\sigma_{Sp}$ |
| --- | --- | --- | --- | --- |
| **ALL** | 87.38% | 0.09 | 88.77% | 0.09 |
| class **A** | 83.31% | 0.12 | 87.48% | 0.06 |
| class **B** | 91.70% | 0.21 | 89.93% | 0.13 |
| class **C** | 98.59% | 0.01 | 98.32% | 0.03 |
| class **D** | 92.01% | 0.06 | 93.43% | 0.03 |
| class **E** | 88.51% | 0.08 | 96.33% | 0.04 |
| class **F** | 65.74% | 0.14 | 69.00% | 0.08 |
| class **G** | 75.66% | 0.15 | 75.35% | 0.13 |

## Data Availability

The repository can be found at https://www.ebi.ac.uk/biostudies/studies/S-BSST81 or http://comprec-lin.iiar.pwr.edu.pl/mirrorModels/.

